# ExPOSE: A comprehensive toolkit to perform expansion microscopy in plant protoplast systems

**DOI:** 10.1101/2024.07.12.603300

**Authors:** Kevin L. Cox, Sarah A. Pardi, Lily O’Connor, Anastasiya Klebanovych, David Huss, Dmitri A. Nusinow, Blake C. Meyers, Kirk J. Czymmek

**Affiliations:** Donald Danforth Plant Science Center, St. Louis, MO 63132; Howard Hughes Medical Institute, Chevy Chase, MD 20815; Plant and Microbial Biosciences Program, Division of Biology and Biomedical Sciences, Washington University in Saint Louis, St. Louis, MO 63130; Division of Plant Science and Technology, University of Missouri, Columbia, MO 65211; The Genome Center, University of California, Davis, Davis, CA 95616; Department of Plant Sciences, University of California, Davis, Davis, CA 95616

## Abstract

Expansion microscopy (ExM) achieves nanoscale imaging by physical expansion of fixed biological tissues embedded in a swellable hydrogel, enhancing the resolution of any optical microscope several-fold. While ExM is commonly used in animal cells and tissues, there are few plant specific protocols. Protoplasts are a widely used cell system across plant species, especially in studying biomolecule localization. Here, we present an approach to achieve robust expansion of plant protoplasts, termed **Ex**pansion microscopy in plant **P**r**O**toplast **S**yst**E**ms (ExPOSE). We demonstrate that coupling ExPOSE with other imaging techniques, immunofluorescence and *in situ* hybridization chain reaction to visualize proteins and mRNAs, respectively, greatly enhances the spatial resolution of endogenous biomolecules. Additionally, in this study, we tested the effectiveness and versatility of this technique to observe biomolecular condensates in *Arabidopsis* protoplasts and transcription factors in maize protoplasts at increased resolution. ExPOSE can be relatively inexpensive, fast, and simple to implement.

## Introduction

The advancement in imaging technologies has led to progress in understanding the structural and molecular organization of cells. Powerful microscopy tools, such as super-resolution microscopy, enable imaging of single molecules and their spatial relationship to other cell and organ structures at nanometer resolution. Viewing these components at an increased resolution has led to new insights into various biological questions (Sydor et al. 2015; Prakash et al. 2022; Czymmek, Duncan, and Berg 2023), particularly in chromatin, RNA, and cell biology (Fornasiero and Opazo 2015). Despite their advantages, super-resolution microscopes are not ubiquitous and require post image processing, and other common optical approaches such as confocal microscopy when used alone still are inherently limited by diffraction.

Expansion microscopy (ExM) can overcome these optical limitations by physically expanding cells and tissues (F. Chen, Tillberg, and Boyden 2015). This isotropic specimen expansion method enables for cost-effective, 3D, nanoscale imaging on even conventional, diffraction-limited microscopes. In ExM, fixed cells and tissues have their molecules covalently anchored to a swellable hydrogel that infiltrates the cells and forms a mesh. Applying water to this gel results in molecules and cellular components physically separating from each other, resulting in a ∼4.5x linear expansion of the specimen (F. Chen, Tillberg, and Boyden 2015). This innovative approach circumvents the resolution limitations of traditional microscopy methods and reveals finer details of cellular structures. ExM is used in various biological applications, including resolving complex subcellular structures, visualizing RNA (ExFISH) (F. Chen et al. 2016) and proteins (proExM) (Tillberg et al. 2016) to understand their organization and sub-structure within organelles and cells, enhancing their localization acuity.

Since its first demonstration in 2015, ExM has been successfully applied across different eukaryotic systems, and numerous variations of the ExM procedure have been produced (Wen et al. 2023). However, the application of this method in the plant kingdom has been limited. Two studies have used ExM in *Arabidopsis thaliana* and these important studies are focused on certain areas of the root (Hawkins et al. 2023) or ovules and seeds with a lower expansion factor (Kao and Nodine 2019, 2021). Outside of *Arabidopsis*, ExM has been applied in the unicellular alga *Chlamydomonas* (Gambarotto et al. 2019; Klena et al. 2023). This general scarcity of plant-specific ExM protocols is primarily due to challenges with their diverse cell walls, which limits uniform penetration of the chemical reagents used for ExM and ultimately restricts cells from expanding. Previous studies have overcome this obstacle by organelle isolation of chloroplasts (Bos, Berentsen, and Wientjes 2023) and nuclei (Kubalová et al. 2020) before applying ExM. However, isolation of these components before ExM can induce deviations to the organelles and miss biological context of the rest of the cell.

For decades, protoplasts have been a highly utilized model for studying cellular processes across different plant species. To date, protoplast techniques have been well established in *Arabidopsis*, maize, rice, wheat, barley, oat, tomato, and many other plant species (Kaur-Sawhney, Flores, and Galston 1980; Kovtun et al. 1998; Takai et al. 2007; Yoo, Cho, and Sheen 2007; Wu et al. 2009; Gomez-Cano, Yang, and Grotewold 2019; Saur et al. 2019; Hahn et al. 2020). Protoplasts are generated by enzymatic digestion of the plant cell wall, making them easily transformable (Cocking 1960). Plant protoplasts provide a high-throughput, versatile system for studying cellular processes such as protein function and localization, signal transduction, transcription regulation, and single-cell multi-omic analyses (Xu et al. 2022). Additionally, these isolated cells allow for individual cell observations with most organelles preserved in their spatial locations (Sheen 2001; Yoo, Cho, and Sheen 2007). These established techniques, combined with maintaining cellular physiology and genetic properties of the whole plant, make protoplasts a useful system to study biological questions.

Thus, given the advantages and versatility of the protoplast system in plant biology research, we set out to develop an ExM protocol leveraging the benefits of single-cell biology with plant protoplasts. This protocol was developed by modifying previously published ExM methods and adapting them for plant protoplasts. We named this method “**Ex**pansion microscopy in plant **P**r**O**toplast **S**yst**E**ms”, or ExPOSE for short. ExPOSE results in the robust physical expansion of whole protoplast cells. We demonstrate the versatility of this method by pairing with other molecular tools to visualize selected proteins and RNA at increased resolution, which is further enhanced via structured illumination based super-resolution microscopy. We show that ExPOSE can also be used to observe biomolecular condensates in *Arabidopsis* protoplasts. Lastly, we revealed the distribution and relationship of two transcription factors in nuclei of maize protoplasts at sub-organelle resolution. This method enhances 3D nanoscale resolution imaging and enables analyses of how proteins and RNAs are spatially organized in plant protoplasts.

## Results

### Establishing an ExPOSE workflow for implementing expansion microscopy in protoplasts

To acquire enhanced imaging resolution and detailed analysis of subcellular components within plant protoplasts, we developed a streamlined ExM protocol termed ExPOSE (Figure 1). First, the cell walls from Arabidopsis leaves and maize etiolated leaves were completely removed via enzymatic digestion to isolate protoplasts. The isolated protoplasts were harvested in 2 mL round-bottom tubes for easier handling during centrifugation and solution exchange. Cells were fixed in paraformaldehyde before being treated with a protein-binding anchor, 6-((acryloyl)amino)hexanoic acid (Acryloyl-X SE), also abbreviated as AcX (Tillberg et al. 2016). Next, the samples were embedded in an active monomer solution that becomes a swellable hydrogel. Cells embedded in the hydrogel are subjected to expansion in water overnight. We found that reverse osmosis (RO) or Milli-Q water gave the best uniform expansion compared to tap water. Other plant-specific ExM protocols often treat the gelled samples with Proteinase K to digest overnight before expansion in water. However, since protoplasts already lack a cell wall, we were able to omit this step with our tested biomolecules. Finally, the cells were imaged to compare the cross-sectional cell size differences of pre- and post-expansion (Figure 2A). The gelled and expanded samples exhibited an average cell area expansion greater than 10-fold (Figure 2B). Furthermore, we stained the protoplasts with Hoechst to compare the DNA architecture inside the nucleus of unexpanded versus expanded cells. The expansion of cells allowed us to observe higher definition of the DNA architecture (Figure 2C, bottom panel). This level of detail was not achievable in unexpanded cells (Figure 2C, top panel). Additionally, ExPOSE was able to discern individual grana within chloroplasts compared to unexpanded cells (compare Fig 2A and 2B, magenta). Together, these results demonstrate the effectiveness of ExPOSE with lattice structured illumination microscopy (SIM) to reliably expand and reveal key sub-organellar features in plant cells, overcoming diffraction limitations.

**Figure 1.**
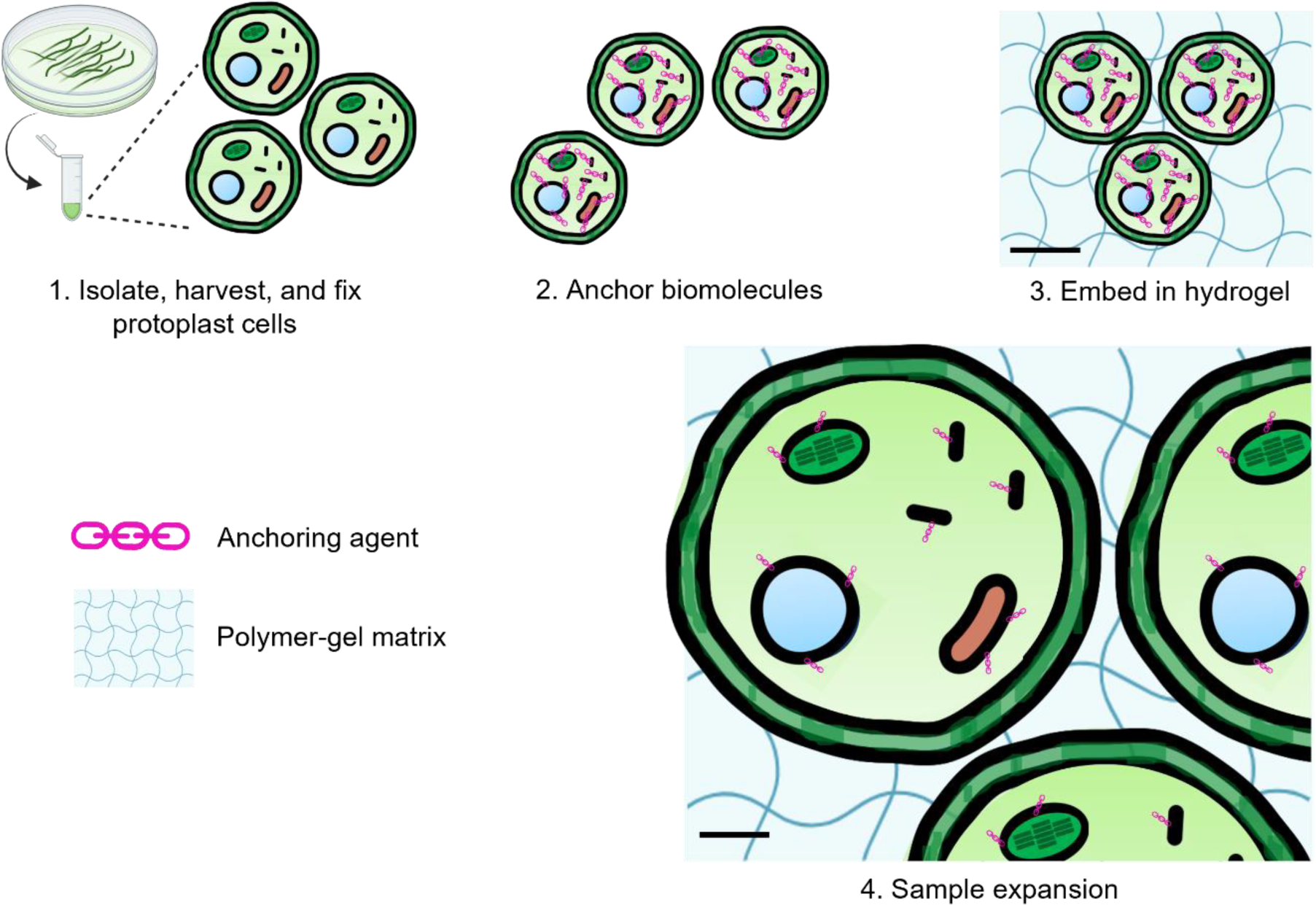
Workflow for expansion of plant protoplast systems (ExPOSE). ExPOSE method for expanding protoplasts by (1) isolating and fixing protoplasts, (2) anchoring the biomolecules, (3) embed the cells in a solution which polymerizes into a gel, and (4) expand the cells embedded in the gel with water, as described in the text underneath each image.

**Figure 2.**
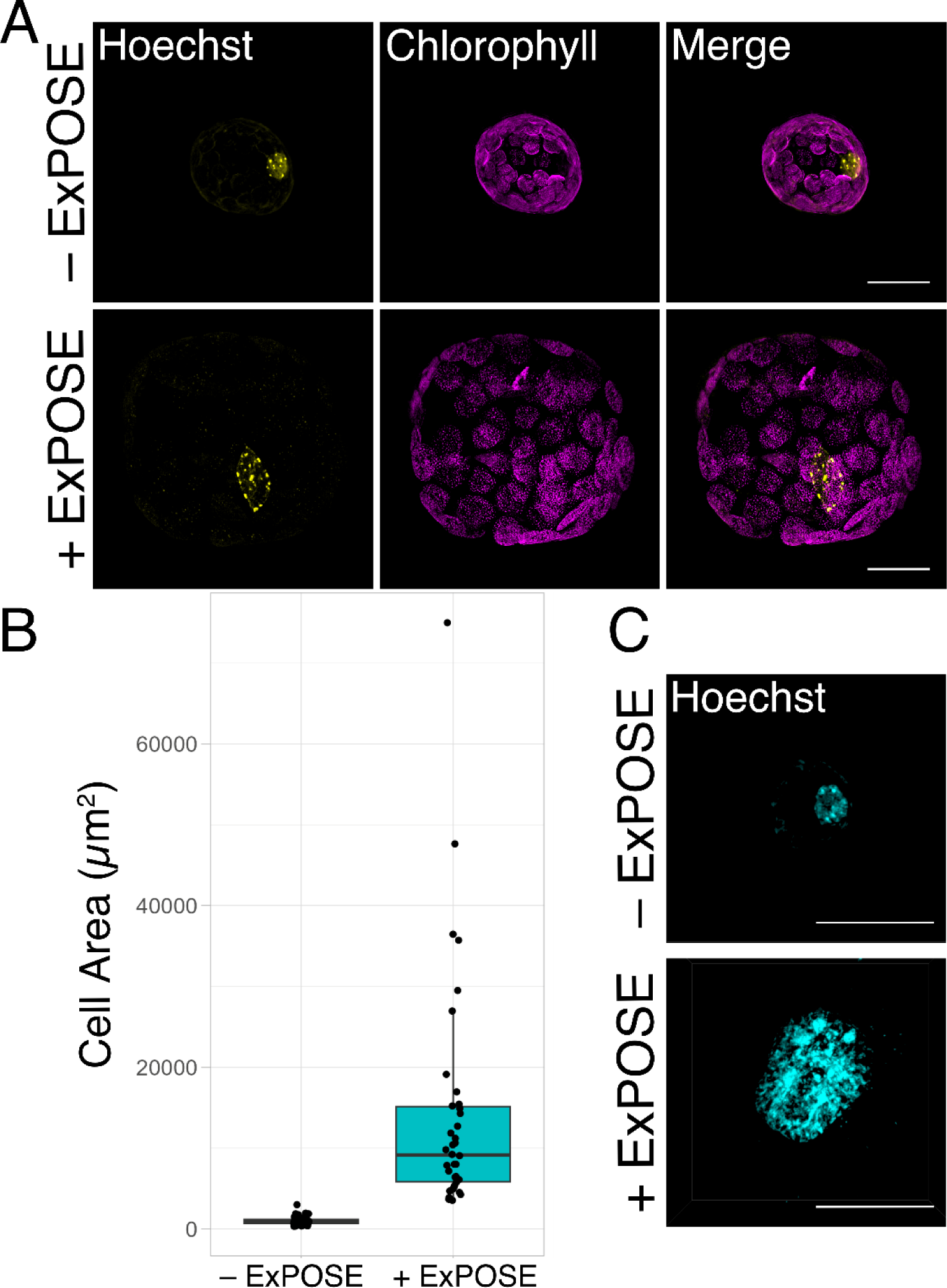
Validation of ExPOSE protocol. (**A**) Lattice SIM maximum intensity projection images of *Arabidopsis* protoplast cells pre- and post-expansion with ExPOSE. DNA stained with Hoechst (yellow), chlorophyll autofluorescence (magenta). Scale bar; 20µm. (**B**) Box plot showing Maximum Intensity Projection (MIP) cell size (area in µm^2^) of unexpanded and expanded protoplast cells (Wilcoxon Rank Sum Test, P = 1.258e-15, n ≥ 38, three independent biological replicates). (**C**) Lattice SIM 3D rendered images of Arabidopsis nuclei pre- and post-expansion. DNA stained with Hoechst (cyan). Scale bar; 20µm.

### ExPOSE enhances the resolution for visualizing endogenous actin and mitochondrial matrix protein localization

After developing the ExPOSE protocol to achieve robust, consistent expansion of protoplasts, we asked whether we can detect endogenous protein localization with the increased resolution afforded by ExPOSE coupled with immunofluorescence and lattice SIM. We applied ExPOSE to *Arabidopsis* protoplasts and labeled actin and mitochondria with anti-actin or anti-mitochondrial matrix (GDC-H: H protein of glycine decarboxylase complex (GDC)) antibodies, respectively. Our post-expanded protoplasts displayed higher definition of both actin and mitochondria localization compared to non-expanded cells (Figure 3A, B). Our ExPOSE method was able to resolve individual actin filaments in expanded protoplasts compared to unexpanded cells (Figure 3A, inset). Furthermore, internal mitochondrial matrices within individual mitochondria were observed, which was not possible to visualize in unexpanded cells, even with lattice SIM (Figure 3B, inset). These results illustrate the potential of coupling ExPOSE with immunofluorescence to enhance visualization of sub-organelle features of endogenous proteins.

**Figure 3.**
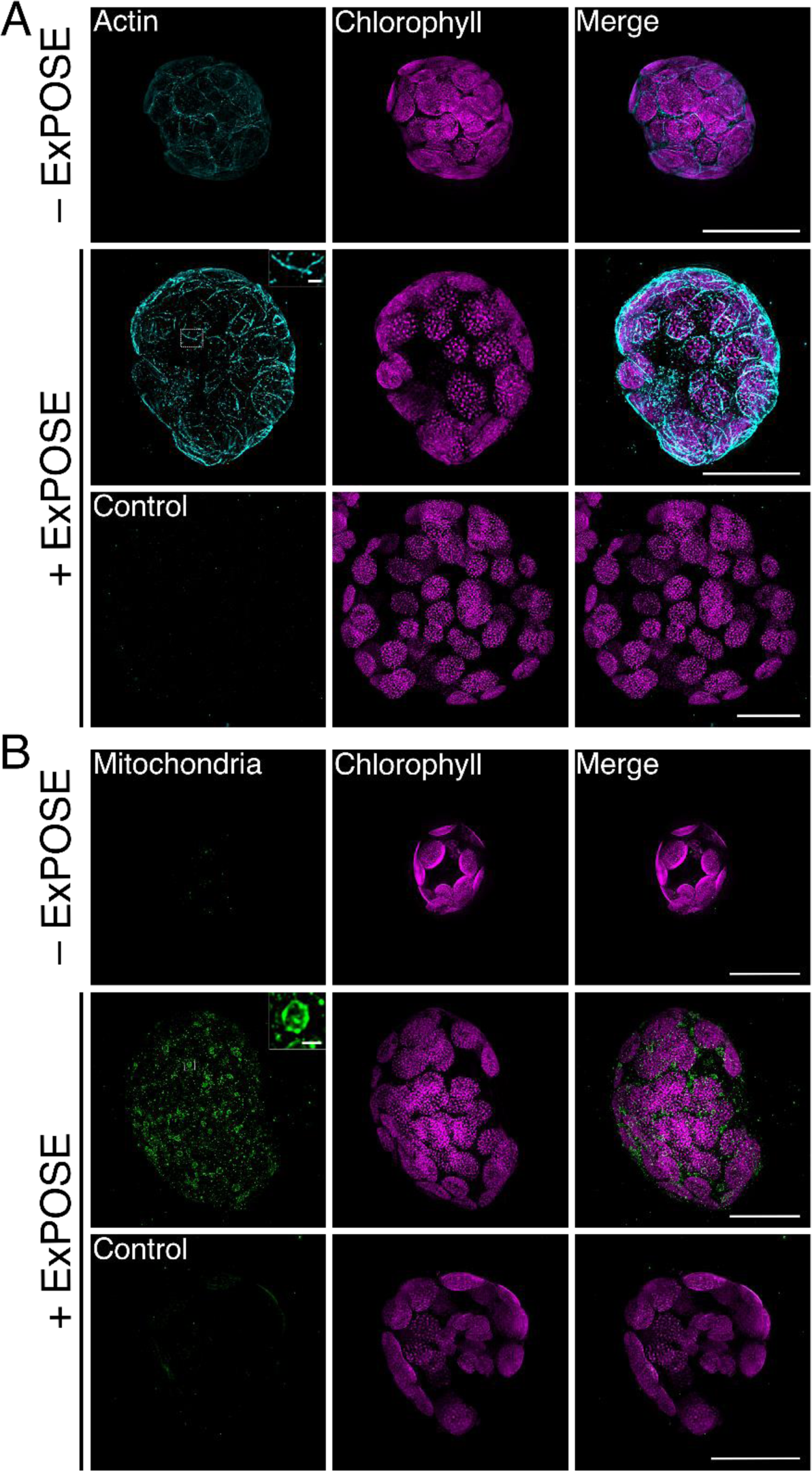
ExPOSE reveals improved resolution of endogenous actin and mitochondria matrix localization in protoplasts. Lattice SIM maximum intensity projection images of *Arabidopsis* protoplasts pre- and post-expansion with ExPOSE and labeled via immunofluorescence using (**A**) anti-actin and (**B**) anti-mitochondrial matrix marker (GDC-H: H protein of glycine decarboxylase complex (GDC)) antibodies, chlorophyll autofluorescence (magenta). Control antibody treatment consisted of incubating samples with non-immune rabbit IgG labeled with DyLight^TM^ 594. Intensity of antibody channels were matched and leveled. Scale bar; 20 µm.

### ExPOSE coupled with in situ HCR enhances detection of individual mRNA foci

After establishing that our ExPOSE method greatly enhanced the spatial resolution of endogenous proteins, we next wanted to test whether our method could enhance the detection and spatial localization of endogenous RNA molecules. To test this, we performed ExPOSE coupled with hybridization chain reaction (HCR). HCR is the targeted hybridization and amplification of a DNA or RNA sequence of interest via hairpin self-assembly cascades (Dirks and Pierce 2004). In this study, we utilized *in situ* HCR v3.0, which fluorescently labels and amplifies target mRNA transcripts while suppressing non-specific background (Choi et al. 2018). We used an anti-sense probe set against *CHLOROPHYLL A/B BINDING PROTEIN 1* (*CAB1*) as our mRNA transcript of interest due to its high abundance in photosynthetic tissues. Our results showed that ExPOSE enhanced signal detection and spatial detail of the labeled *CAB1* mRNA foci compared to non-expanded cells (Figure 4). Our non-specific amplification (NSA) control of the fluorescent hairpins displayed no mRNA labeled amplification, as expected (Figure 4, bottom row). Overall, our results demonstrate the high sensitivity and magnification that ExPOSE, in combination with lattice SIM, provides in revealing the fine detail of individual mRNA foci localization in single-cell protoplasts.

**Figure 4.**
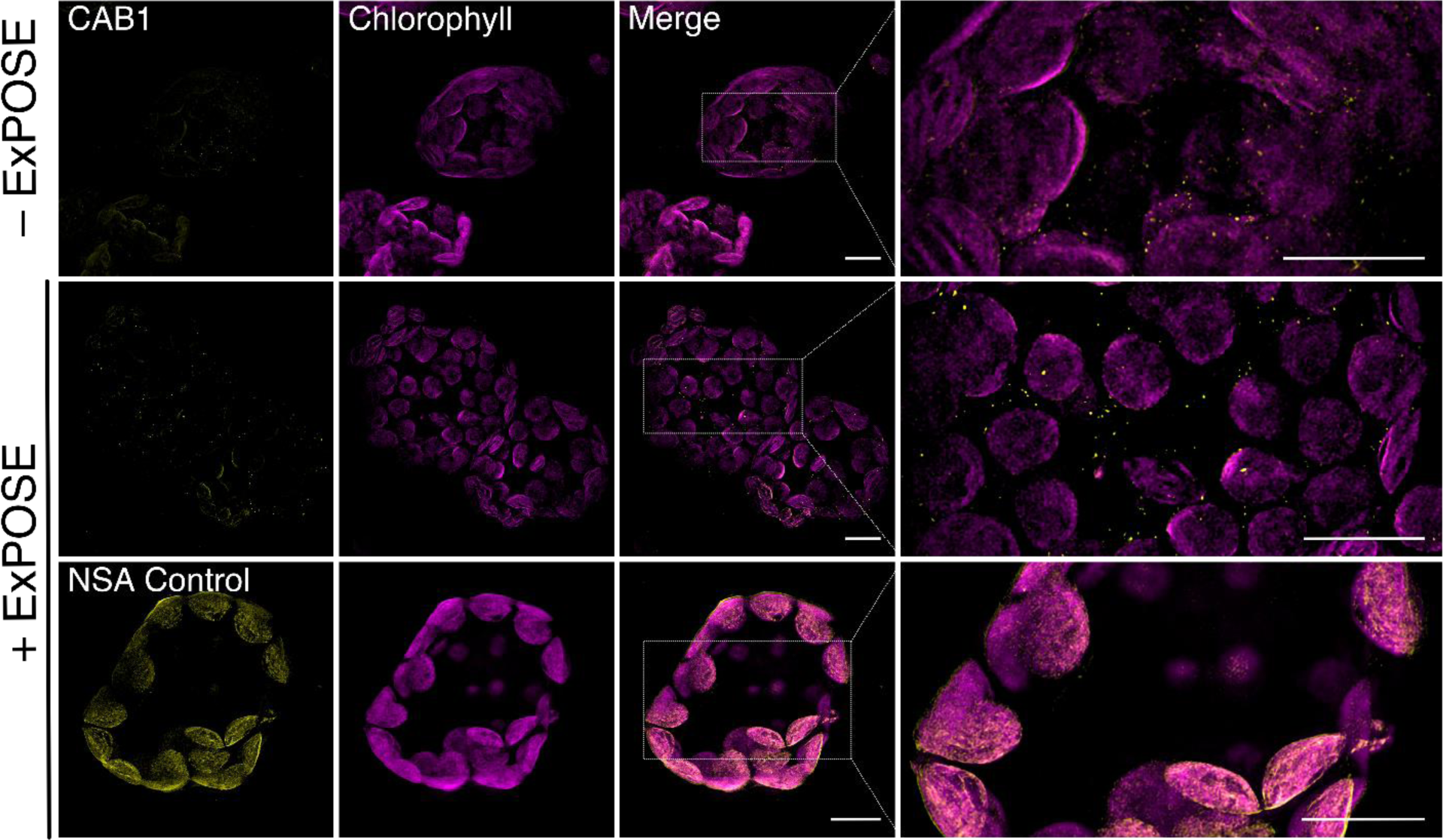
ExPOSE enhances detection of *CAB1* mRNA foci in protoplasts. Lattice SIM maximum intensity projection images of *CAB1* mRNA labeled using HCR (Alexa Fluor^TM^ 546, yellow puncta) in *Arabidopsis* protoplasts pre- and post-expansion via ExPOSE, chlorophyll autofluorescence (magenta). A non-specific amplification (NSA) control is shown in which only fluorescently tagged hairpins were used. Scale bar; 10 µm.

*ExPOSE can be used to visualize biomolecular condensates in Arabidopsis protoplasts* Biomolecular condensates in the field of plant cell biology have gained much attention over the last decade, as they act as cellular sensors to the outside environment (reviewed in Emenecker, Holehouse, and Strader 2021; Field et al. 2023). They are characterized as membraneless subcellular compartments consisting of proteins and nucleic acids (Hyman and Brangwynne 2011), and serve many different cellular functions, including transcription regulation, RNA processing, protein homeostasis, macromolecule storage, and signal transduction (Banani et al. 2017). Different microscopy tools have been used to characterize condensate morphology and substructure. However, expansion microscopy has yet to be performed to investigate biomolecular condensates. Since condensates are often dynamic and transient structures, we next wanted to test whether ExPOSE can maintain the integrity of and be used to visualize biomolecular condensates. For this study, we chose to utilize *Arabidopsis* Phytochrome B (phyB) photobodies as our condensate of interest, which functions as a photosensor and thermal sensor (Sharrock and Quail 1989; Jung et al. 2016; Legris et al. 2016). Phytochromes are red / far-red light-sensing photoreceptors found across kingdoms, in plants, fungi, and bacteria (Mathews and Sharrock 1997; Buchberger and Lamparter 2015). Upon activation via red-light, phyB undergoes phase separation and forms nuclear condensates called photobodies (Yamaguchi et al. 1999; D. Chen et al. 2022). To test whether ExPOSE can be used to observe photobodies, we isolated protoplasts from the stable transgenic *Arabidopsis* 35S::PhyB-GFP line, exposed the cells to red light to stimulate photobody formation, and performed our ExPOSE method. Upon observation, our results showed that ExPOSE preserved phyB-photobody morphology (Figure 5). The physical cell expansion by ExPOSE combined with lattice SIM aided in the spatial detection of individual condensates while decreasing overcrowded obstruction by other cellular organelles, like chloroplasts (Figure 5). This demonstrated that our ExPOSE method can be leveraged to enhance our ability to study biomolecular condensate native structure in single-cell plant protoplasts.

**Figure 5.**
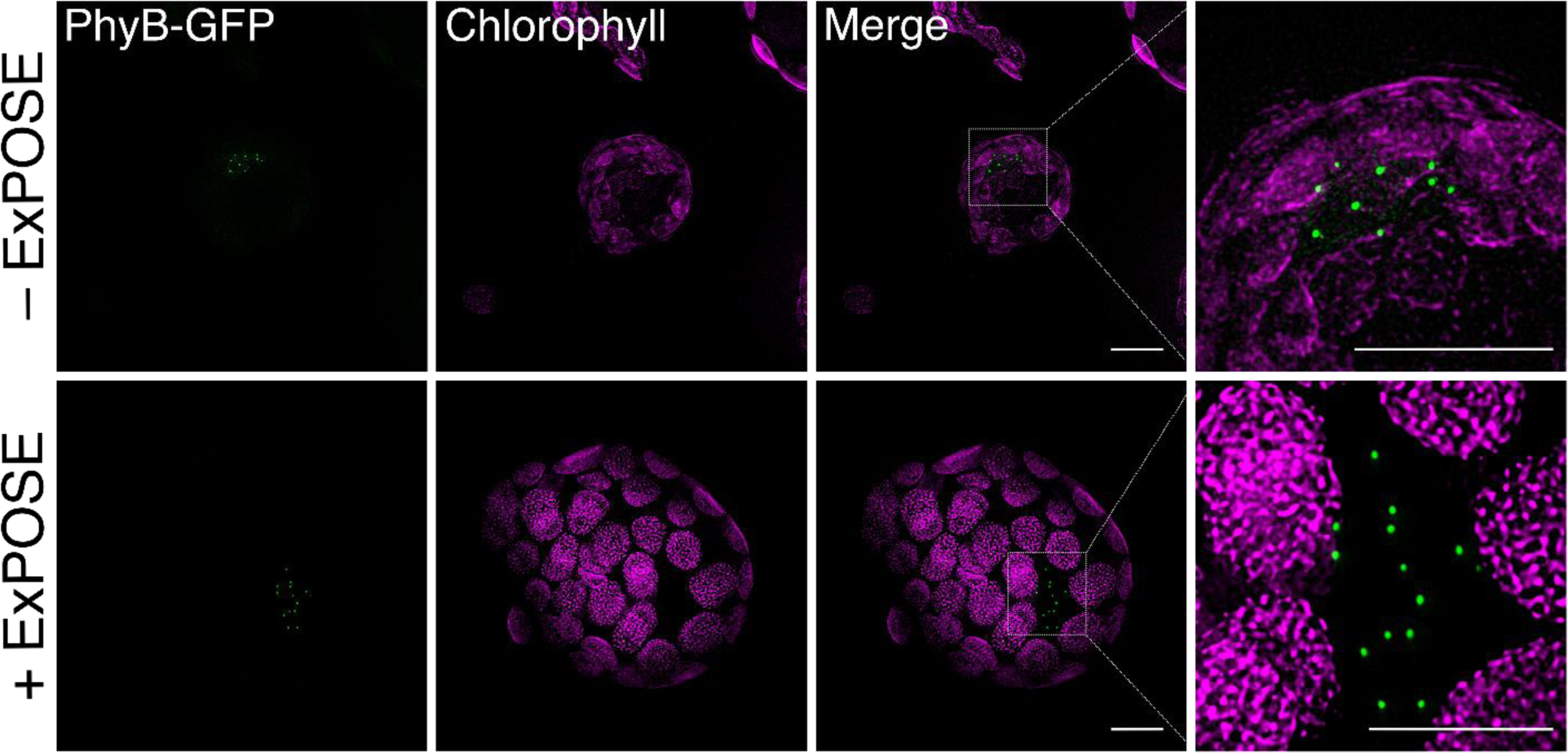
ExPOSE can be used to image biomolecular condensates in protoplasts. Lattice SIM maximum intensity projection images of pre- and post-expanded 35S::PhyB-GFP (green) stable transgenic *Arabidopsis* protoplasts, chlorophyll autofluorescence (magenta). Cells were treated with 10 µmol/m^2^/s red-light to stimulate photobody formation before undergoing ExPOSE. Scale bar; 10 µm.

### ExPOSE reveals the localization patterns of two basic helix-loop-helix transcription factors in maize protoplasts

ExM enables improved visualization of subcellular components within organelles, such as proteins in the nucleus. This allows closely packed proteins, such as transcription factors, to be visualized at greater spatial resolution, potentially providing further insights into their distribution and functions. We applied ExPOSE to maize (*Zea mays*) leaf protoplasts to observe the localization of two maize basic helix-loop-helix (bHLH) transcription factors, *Male Sterile* (*MS*) *23* and *MS32* (Chaubal et al. 2000). MS23 and MS32 are required for maize anther fertility, and yeast two-hybrid and protoplast data suggests that they physically interact to form a heterodimer (Chaubal et al. 2000; Moon et al. 2013; Nan et al. 2017; Nan et al. 2022). To gain further insights into how these two bHLH transcription factors organize and function within maize cells, we ectopically expressed fluorescently tagged MS23 and MS32 in maize leaf protoplasts (p35S:MS23-GFP or p35S:MS32-mCherry) and subsequently performed ExPOSE. After expansion, we stained DNA with Hoechst and observed the localization of these two transcription factors with lattice SIM. ExPOSE further resolved the localization of MS23 (Figure 6A) and MS32 (Figure 6B), and demonstrated that they have unique localization patterns, especially within the nucleus. MS23 localized almost exclusively to the nucleus, where it is distributed in regions with less densely packed DNA, as indicated by Hoechst staining (Figure 6A). While MS32 has both nuclear and cytoplasmic localization, its localization within the nucleus was primarily sequestered to a single concentrated region (Figure 6B). Next, we co-transfected both p35S:MS23-GFP and p35S:MS32-mCherry into maize leaf protoplasts and performed ExPOSE to visualize their distribution using confocal microscopy (Figure 6E) and lattice SIM (Figure 6C). MS23 and MS32 exhibited different localization patterns when co-transfected compared to when transfected individually. In cells co-expressing MS23 and MS32, the fluorescent signal of MS23 becomes more uniformly distributed throughout the nucleus (Figure 6A, 6C, 6D & 6E). When co-transfected with MS23, MS32 cytoplasmic localization decreased, and its nuclear localization becomes more diffuse (Figure 6B-E). When co-transfected, the changes in the nuclear localization of MS23 and MS32 are more readily observed in cells where ExPOSE was performed, especially in the case of MS23 where localization changes are more subtle. In co-transfected cells, the change in nuclear localization of MS23 was only readily observable in unexpanded cells when lattice SIM was used. However, this change was easily observed using confocal microscopy in ExPOSE processed cells and lattice SIM images of ExPOSE samples add even greater acuity. The ExPOSE cells demonstrate that MS23 and MS32 co-expression results in distinct spatial organization of both MS23 and MS32 within maize nuclei when compared to MS23 or MS32 expressed alone. These results demonstrated the utility of ExPOSE for visualizing cellular proteins that are localized in subcellular organelles at an increased resolution and coupling ExPOSE with lattice SIM yields new insights on how these proteins may interact.

**Figure 6.**
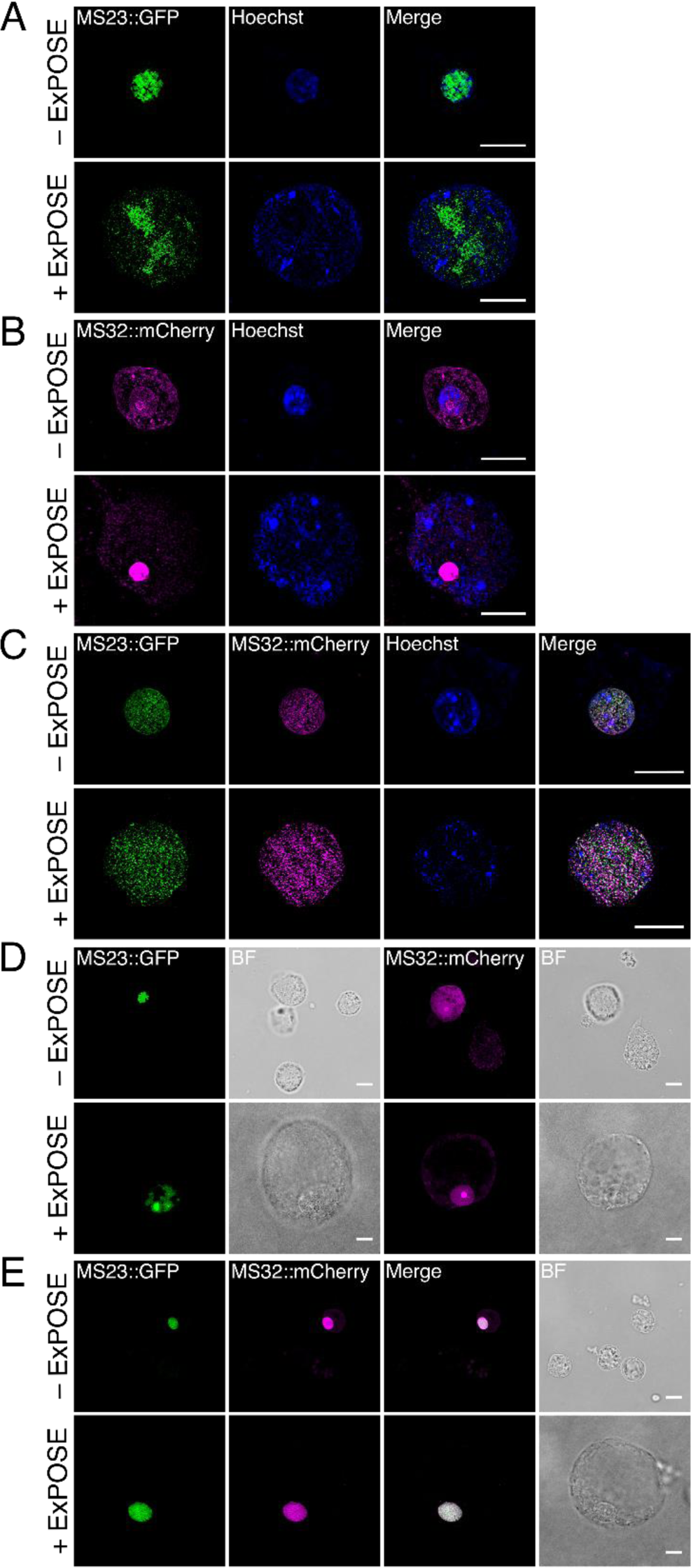
ExPOSE reveals the localization pattern of two basic helix-loop-helix transcription factors when expressed alone versus together in maize protoplasts. **(A-C)** Lattice SIM maximum intensity projection images of pre- and post-expanded maize protoplast cells transiently transfected with (**A**) MS23-GFP (green) alone, (**B**) MS32-mCherry (magenta) alone, and (**C**) MS23-GFP and MS32-mCherry together. DNA stained with Hoechst (blue). (**D-E**) Deconvolved maximum intensity projection confocal images of pre- and post-expanded maize protoplast cells transiently transfected with either (**D**) MS23-GFP (green) alone, or MS32-mCherry (magenta) alone, or (**E**) MS23-GFP and MS32-mCherry together. BF = Brightfield. Scale bar; 10 µm.

## Discussion

In plant research, protoplasts are a widely used transient cell system across monocots and dicots (Xu et al. 2022). Protoplast transformation is a rapid way to test protein localization, protein-protein interactions, transcription regulation, among other cellular processes, eliminating the length of time needed to generate stable transgenic lines. With the recent advancements in plant imaging to uncover the structural and molecular organization of biomolecules (Ovečka et al. 2021), protoplasts provide a unique single-cell system to study cellular and subcellular properties. Here, we developed the method ExPOSE as a tool to reliably expand plant protoplasts to visualize subcellular components. Cells are crowded environments, so this physical expansion method allows more detailed observations of biomolecules of interest by simply increasing the relative cell volume by several fold. ExPOSE, and other ExM techniques, not only circumvent the inherent optical diffraction limit for a given objective lens and imaging modality but can be further enhanced in combination with super-resolution microscopy, such as lattice SIM as applied here. This protocol provides robust expansion of cells, is performed in a span of a few days, is minimally labor intensive, and can be performed in any plant biology laboratory.

Once protoplasts are generated, this ExM method has the advantage of being applicable to different plant systems with few modifications. Generally, due to the plant cell wall, when a method is developed for one plant species, extensive time is spent optimizing that same method to apply to a different species (e.g., plant transformation, transient silencing of genes, nuclei/protein isolation). While protoplast isolation techniques vary across plant species, the downstream protocol to perform ExPOSE after isolation is easily transferable, as demonstrated by using Arabidopsis and maize protoplasts in our study. On the other hand, it is noted that there are plant systems for which protoplast isolation is not achievable or efficient. However, the rise of plant single-cell RNA-sequencing studies has fueled the optimization of protoplast isolation protocols from non-model species (Zheng et al. 2023). This holds promise for making ExPOSE applicable and complementary to single-cell studies for many more plant species in the near future.

Immunofluorescence and HCR are excellent molecular tools for visualizing the localization patterns of endogenous proteins and nucleic acids, respectively, bypassing the need for creating fusion proteins and cell transformation. Coupling those tools with lattice SIM and ExPOSE, as we showed here, greatly improved visualization of individual actin filaments and the mitochondrial matrix, which was not possible without cell expansion. Furthermore, we were able to visualize transcripts of *CAB1* mRNA as individual or clustered foci. Thus, coupling molecular methods with ExPOSE is a powerful toolkit for high-resolution biomolecule imaging in plant cells.

Additionally, ExPOSE can be used for imaging biomolecular condensates with greater spatial resolution. The form and function of biomolecular condensates in plant systems have garnered much attention over the last decade. There are numerous examples of plant nuclear and cytoplasmic bodies, including Cajal bodies, stress granules, Auxin Response Factor (ARF), Flowering Control Locus A (FCA), Nonexpressor of Pathogenesis-Related Genes 1 (NPR1), EARLY FLOWERING 3 (ELF3) condensates, among others (reviewed in Emenecker, Holehouse, and Strader 2020; Field et al. 2023). Here, ExPOSE was applied to determine suitability for preserving native spatial localization of condensates, specifically of phyB photobodies. With many types of condensates present in cells, this method can be used to reveal overlapping or segregating condensate spatial positioning and functions. Some biomolecular condensates are highly complex structures, displaying diverse architectures and even containing sub-compartments, such as inner core - outer shell phenotypes (Fare et al. 2021). Additionally, different proteins or RNAs can localize to different sub-compartments (King, Ruff, and Pappu 2024). Since biomolecular condensates are made up of proteins and often nucleic acids, ExM can serve as an excellent tool to observe their subcompartments with improved clarity, including RNA distribution within RNA containing condensates. Our ExPOSE approach is straightforward, and can be complementary to other, more expensive and labor intensive high resolution imaging techniques, such as single-molecule localization microscopy (SMLM) and transmission electron microscopy (TEM), used to achieve nanometer resolution for uncovering condensate architecture (Pandey, Budhathoki, and Spille 2023; Ibrahim et al. 2024). Here, we demonstrated that ExPOSE in combination with lattice SIM worked effectively for visualization of condensates, and thus can be useful for studying condensate morphology with enhanced resolution in plant protoplasts.

The spatial organization and interactions of proteins within the nucleus are complex and can often be difficult to resolve. Here, we show that ExPOSE is a useful tool for investigating the spatial localization of transcription factors in protoplasts. In maize protoplasts transiently expressing two bHLH transcription factors, MS23 and MS32, ExPOSE revealed that the spatial organization of MS23 and MS32 within maize nuclei is regulated by the presence of both transcription factors. The enhanced resolution of the ExPOSE results allowed for subtle changes in nuclear localization to be observed. Many eukaryotic transcription factors form dimers with similar or identical molecules and bind to DNA sequences at a much higher specificity (Amoutzias et al. 2008). ExPOSE provided a way to resolve these localizations at much higher resolution in a relatively rapid workflow. While we utilized ExPOSE to investigate transcription factor:transcription factor spatial localization patterns, this method could be applied to investigate other nuclear localization patterns, including DNA:protein or RNA:protein localization. ExPOSE has the potential to reveal nuclear organization under various transcriptional, environmental, or developmental states, which can provide additional information on chromatin:chromatin relationships along-side methods such as high-throughput chromosome conformation capture (Hi-C). Additionally, chromatin expansion microscopy (ChromExM), has recently been developed in zebrafish embryos for understanding nascent transcription (Pownall et al. 2023).

ExM methods for plant samples are still in their infancy. Here, we present ExPOSE as a robust approach for expanding plant protoplasts for imaging single-cell and subcellular compartments at greatly enhanced resolution. ExM of intact organs and whole organisms will be the next target of interest in plant biology, as this would provide more tissue level context and 3D spatial information of cellular organelles and other biological questions. Recently, studies have reported the use of ExM on intact *Arabidopsis* roots, ROOT-ExM, to achieve expansion to a factor of 4x along with super-resolution imaging (Gallei et al. 2024; Grison et al. 2024). As technical issues are surmounted, the increasing momentum of this field will see the rise of more accessible plant-specific ExM to visualize subcellular components and their spatial relationships during cell division, biotic/abiotic interactions, and other biological studies.

## Experimental Procedures

### Plant materials and growth conditions

*Arabidopsis thaliana* accession Col-0 (WT) and 35S::PhyB-GFP (Huang et al. 2016) seeds were sterilized, plated on ½ MS 1% Sucrose, and stratified in the dark at 4°C for two days. Plates were placed in chambers with a photoperiod of 12 hr light:12 hr dark at 22°C. After two weeks, seedlings were transferred from plates to soil to grow in long day conditions (16 hr light:8 hr dark) with 50% relative humidity for another two weeks. Leaves from 4-week old plants were collected for protoplast isolation.

For maize, seeds of B73 were germinated in soil and grown in constant darkness at 25°C and 50–70% relative humidity. Leaves were collected from etiolated seedlings 10–12 days after germination for protoplast isolation.

### Protoplast isolation and transfection

*Arabidopsis* protoplast isolation was performed as previously described, with slight modifications (Hansen and van Ooijen 2016). Briefly, 4-week old *Arabidopsis* leaves were cut, mesophyll cells were exposed via the tape-sandwich method, and protoplasts were released in enzyme solution (0.5% w/v Cellulase, 0.25% w/v Maceroenzyme, 400 mM D-mannitol, 10 mM CaCl_2_, 20 mM KCl, 0.1% w/v BSA, 20 mM MES pH 5.7). Protoplasts were collected via centrifugation (100 x g for 3 min at 4°C), resuspended in W5 solution (150 mM NaCl, 125 mM CaCl_2_, 5 mM KCl, 2 mM MES pH 5.7), rested on ice in the dark for 30 minutes, centrifuged again, and finally resuspended in W5 with 3.2% paraformaldehyde. Before fixation for PhyB-GFP, protoplasts were exposed to red light (10 µmol/m^2^/s) at room temperature for 30 minutes to promote formation of PhyB photobodies.

Maize leaf protoplast isolation was performed as previously described (Gomez-Cano, Yang, and Grotewold 2019). Briefly, the second leaf of 12-day old dark grown B73 seedlings were sliced into ∼1 mm sections, submerged in protoplast enzyme solution (0.6 M mannitol, 20 mM KCl, 20 mM MES pH5.7, 10 mM CaCl_2_, 0.1% w/v BSA, 1.5% w/v cellulase “ONOZUKA” RS, 0.4% w/v Macerozyme R-10), and placed under a vacuum for 40 mins. The vacuum was removed and digestion continued for 3 hrs with rotation (60 RPM) at room temperature. Digestion was halted by adding an equal volume of W5 solution and tissue debris was removed by filtering solution through a 40 µm filter. The isolated protoplasts were collected by centrifugation (100 x g 3 min) and resuspended in W5 solution. Protoplasts were rested on ice for about 30 minutes and then resuspended in room temperature MMG solution (0.6 M mannitol, 15 mM MgCl_2_, 4 mM MES pH 5.7) to a final concentration of 2.5 x 10^6^ cells/mL.

For maize protoplast transfections, 3 pmols of plasmid DNA was added to 2.5 x 10^5^ protoplasts in MMG and an equal volume of PEG transfection solution (40% PEG4000, 0.3 M mannitol, 0.1 M CaCl_2_) was added. After incubating for 5 minutes at room temperature, transfection was stopped by adding 2x volume of W5. Protoplasts were collected (100 x g 3 min) and stored in WI solution (0.6 M mannitol, 4 mM MES pH 5.7, 20 mM KCl) overnight at room temperature in the dark. Plasmid DNA for transfections was prepared using the ZymoPURE II Plasmid Midiprep Kit (Zymo Research). The p35S:GWC-GFP vector, MS23 coding sequence in pENTR, and MS32 coding sequence in pENTR were previously utilized by Nan et al., 2022 (Nan et al. 2022). Briefly, the MS23 and MS32 coding sequences were obtained from the Maize TFome collection and the p35S:GWC-GFP backbone vector was a gift from Professor Erich Grotewold. The p35S-GWC-mCherry backbone vector was generated by rearrangement of the original backbone via restriction digestion cloning, followed by insertion of mCherry coding sequence via NEBuilder® HiFi DNA Assembly. The final p35S:MS23-GFP and p35S:MS32-mCherry expression vectors were obtained using Gateway LR Clonase recombination (ThermoFisher).

After isolation and transfection, protoplasts were harvested in 2 mL round-bottom microcentrifuge tubes and resuspended in 3.2% paraformaldehyde in WI (maize) or W5 (Arabidopsis) buffer and fixed overnight at 4°C.

### Preparation of AcX and LabelX reagents

“AcX” (6-((acryloyl)amino)hexanoic acid, succinimidyl ester; also known as Acryloyl-X, SE; Thermo Fisher Scientific) and “LabelX” reagents were prepared as previously described (Asano et al. 2018; Zhang et al. 2020). Briefly, stock solutions of AcX were prepared at a final concentration of 10 mg/ml in DMSO. The LabelX reagent was prepared by firstly resuspending Label-IT Amine Modifying Reagent (Mirus Bio, LLC) was resuspended in the provided Reconstitution Solution at 1 mg/mL, followed by reacting 100 uL of Label-IT Amine Modifying Reagent stock solution (at 1 mg/mL) to 10 uL of AcX stock solution overnight at room temperature. AcX and LabelX aliquots were stored at -20°C in a sealed container with a desiccant (Drierite).

### AcX and LabelX treatment

Fixed protoplasts were washed three times by adding WI and W5 buffer to maize and *Arabidopsis* protoplasts, respectively, and resuspending cells. For treatment, protoplasts were resuspended in 0.01 mg/mL of AcX (for normal ExM) or LabelX (for HCR-ExM) in WI (maize) or W5 (*Arabidopsis*) buffer and incubated in the dark, for overnight at room temperature. After the overnight incubation, samples were washed three times with their corresponding buffer (WI or W5) before proceeding with gelation and expansion.

### Gelation and expansion for ExM in protoplasts

A monomer solution (“Stock X’) made of 8.6% (w/v) sodium acrylate, 2.5% (w/v) acrylamide, 0.15% (w/v) N,N-methylenebisacrylamide, 2 M sodium chloride, and 1x PBS was prepared, aliquoted, stored at -20°C, and thawed before use. Immediately before gelation, tetramethylethylenediamine (TEMED) and ammonium persulfate (APS) were added to an aliquot of Stock X at a final concentration of 0.1% (w/v) TEMED and 0.1% (w/v) APS. The solution was briefly vortexed and placed on ice to prevent premature polymerization. After pelleting the protoplasts and removing the buffer, 50 uL of the gelation solution was added to the samples before immediately transferring them to silicone incubation chambers (Electron Microscopy Sciences, Cat. No. 70324-05) and sealing with a glass microscope slide. Slides were incubated for 1 hour at 37°C on a rotisserie rotator in a hybridization oven, a critical step to complete homogeneous cell distribution and polymerization of within the gels. Carefully, a razor blade was used to slide between the silicon mold and microscope slide to release the polymerized gel into a container with 30 mL RO or Milli Q water. The gels were allowed to expand in water overnight, shaking at 60 RPM at room temperature in the dark.

### Hybridization chain reaction on expansion microscopy samples

Oligonucleotide probe sets for HCR were designed against *Arabidopsis thaliana CAB1* (*chlorophyll A/B binding protein 1*, AT1G29930) mRNA transcripts (Choi et al. 2018). 15mm filter inserts (Netwell Insert, 74 µm polyester mesh, Costar) were added to the wells of a standard 12 well tissue culture dish. Each protoplast gel was scraped from the glass slide into a single well containing 5x SSCT (750 mM sodium chloride, 0.75 mM trisodium citrate with 0.1% Tween-20). The filter inserts were then moved into wells containing 3 mL of hybridization buffer (30% formamide, 5x SSC, 9 mM citric acid, 1X Denhardt’s solution, 10% dextran sulfate, 0.1% Tween-20) and incubated at 37°C for 30 min. The probe sets were diluted in hybridization buffer at a final concentration of 20 nM/probe. Two wells per experiment received no probes. The gels were incubated at 37°C overnight with slight agitation. The gels were washed twice in 15% formamide, 5X SSC, 4.5 mM citric acid, 0.1% Tween-20 at 37°C for 15 min, followed by 2x SSCT (300 mM sodium chloride, 30 mM trisodium citrate, 0.1% Tween-20) at 37°C for 15 min and a final 15 min wash in 5x SSCT at RT. After washing, the gels were transferred into 1.5 ml microfuge tubes with fresh 5x SSCT.

A pair of Alexa Fluor^TM^ 546 conjugated hairpins (H1 and H2, Molecular Instruments, Los Angeles, CA) corresponding to the amplifier sequence of the probe sets were pipetted into separate microfuge tubes and heated to 95°C for 90 sec in a heat block, then allowed to refold at RT for 30 min in the dark. The tubes were then spun down and the hairpins added to the amplification buffer (5x SSCT, 10% dextran sulfate) at a final concentration of 60 nM. The 5x SSCT was replaced with the hairpin amplification buffer and incubated overnight at room temperature in the dark. One gel per experiment received no hairpins. The hairpin buffer was removed and the gels washed 3 x 10 min in 5x SCCT at RT. The gels were then expanded as above.

### Post-expansion immunostaining

For immunofluorescence staining, the samples were processed either as AcX-treated isolated protoplasts in suspension (referred to as ExPOSE(-)) or as unexpanded, reduced-size gels which were subsequently expanded (referred to as ExPOSE(+)) in RO water after final washing steps. Prior to incubation with a primary antibody, samples were incubated in a blocking buffer containing 2% (w/v) BSA in 1X PBS buffer pH 7.4 for one hour at RT on a shaker. The primary antibodies used in this study were anti-actin rabbit polyclonal (AS13 2640, Agrisera, Vannas, Sweden), and mitochondrial matrix marker anti-GDC-H rabbit polyclonal antibody (AS05 074, Agrisera, Vannas, Sweden) at a dilution of 1:250 in blocking buffer. The samples were incubated with the diluted primary antibodies at 4°C on a shaker overnight. Following the incubation, the gels were washed three times in a blocking buffer for 10 min each on a shaker at room temperature. Similarly, the protoplasts in suspension were briefly centrifuged, the supernatant was removed with a wide-bore tip, and three consecutive washing steps with centrifugation were performed as described above. After washing steps, the samples were incubated with the secondary donkey anti-rabbit antibody conjugated with DyLight^TM^ 594 (AS12 2076, Agrisera, Vannas, Sweden), diluted in a blocking buffer to a final concentration of 1:500. We allowed the samples to incubate for 3 hours on a shaker at room temperature, followed by three consecutive washing steps in 1X PBS buffer as mentioned above. Finally, the gels were left to expand in RO water overnight on a shaker at RT in the dark.

### Imaging and analysis

Before imaging, the protoplast suspension was mounted on a microscope slide (Cat No. 71883-05, Electron Microscopy Sciences, Hatfield, PA, USA) using Secure Seal™ imaging spacer (Cat No. 70327-9S, Electron Microscopy Sciences, Hatfield, PA, USA) and square cover glass No. 1.5 (Cat No. 722204-01, Electron Microscopy Sciences, Hatfield, PA, USA). Expanded and unexpanded gels were imaged using high-precision glass-bottom dishes (Cat No. HBSB-5040, WillCo Wells, Amsterdam, The Netherlands) treated with 0.1% (w/v) poly-L-lysine solution (Cat No. P8920, Sigma-Aldrich, USA) according to the manufacturer’s protocol with minor modifications. Briefly, we covered the surface of the glass-bottom dishes with 0.1% poly-L-lysine water solution for about an hour at RT, followed by a quick water rinse and subsequent overnight heating step on a hot plate set to 60°C. Laser scanning confocal microscope images were taken with a Leica TCS SP8 with white light lasers using the Leica Las X software and the following objectives: 63x/1.2 HC PL APO CS2 water immersion, 40x/1.1 PL APO CS2 water immersion and 20x/0.7 HC PL APO air immersion. Images were taken at 1024 x 1024 pixels with a bit depth of 12. Hoechst 3342 was excited using the 405 nm diode laser line and emission was collected at 415-475 nm. Chlorophyll was excited using the 633 nm laser line and collected at 643-750 nm. Lattice SIM super-resolution images were taken with a ZEISS Elyra 7 inverted microscope running the Zen Black 3.0 SR software package and a 40x/NA 1.2 C-Apochromat Corr FCS water immersion objective and a 63x/NA 1.2 C-Apochromat M27 water immersion objective. Consecutive z-stack images were taken at 1024x1024 or 1280×1280 pixel frame size and at 16-bit depth. Hoechst 33342 was excited with 405-nm; Chlorophyll a/b, PhyB-GFP and *CAB1* with 642-nm, 488-hm and 561-nm lasers, respectively. Multi-channel imaging was done sequentially in fast frame mode. Respective emissions were detected using the pco.edge sCMOS camera and BP 420-480 + BP 495-550 + LP655 filter or BP 495-550 + BP 570-620 or BP 570-620 + LP 655 filter. The ZEISS SIM^2^ processing module was used to reconstruct SIM super-resolution images, followed by a deconvolution and option “scale to raw image” was selected to retain original relative signal intensities. Experimental and control images were taken using identical microscope settings as described above. For visualization of expansion, the brightness levels were adjusted for qualitative comparison of size and morphology but not the intensity of the signals. In Fig. 3 A-B, left panel images showing fluorescent antibody labeling were displayed with matching black and white levels so that the intensity of the signal was leveled. All images were processed to obtain Maximum Intensity Projections (MIPs) and exported to 8-bit RGB TIFF format for final figure assembly.

### Statistical analysis

Statistical analyses were conducted using R version 4.3.2 (R Foundation for Statistical Computing 2018). Shapiro-Wilk normality test showed that data deviate from a normal distribution. The Wilcoxon Rank-Sum test was used to determine whether two independent populations were significantly different. The statistical analysis was conducted on a total sample size of 38 for expanded protoplasts and on a total sample size of 50 for unexpanded protoplasts, comprising 3 individual biological repeats.

### Analysis of expansion area

Area measurements were performed in Fiji open-source software (Schindelin et al. 2012). The images acquired as Z-stacks with chlorophyll a/b were used as a reference channel to outline the protoplast perimeter. To construct a 2D image for subsequent analysis, individual Z stacks were taken and applied the Maximum Intensity Projection (MIP) algorithm. Finally, the MIP area in μm^2^ for individual protoplasts was calculated.

## Acknowledgements

This work was supported by the Howard Hughes Medical Institute (Hanna Gray Fellowship award to K.L.C.), the U.S. National Science Foundation (award #1945854 to B.C.M.), Washington University in Saint Louis (William H. Danforth Fellowship in Plant Sciences awarded to L.O.), and the William H. Danforth Fellowship (awarded to S.A.P). We also gratefully thank the NSF for partial funding this work (Award # 2130365 to K. Czymmek) and Joanna Porankiewicz-Asplund and Agrisera for providing anti-actin, mitochondrial matrix primary antibodies and DyLight^TM^ secondary antibodies. Imaging support was provided by the Advanced Bioimaging Laboratory (RRID:SCR_018951) at the Danforth Plant Science Center and usage of the ZEISS Elyra 7 Super-Resolution Microscope acquired through an NSF Major Research Instrumentation grant (DBI-2018962) and Leica SP8-X confocal acquired through an NSF Major Research Instrumentation grant (DBI-1337680).

## Author Contributions

K.L.C., S.A.P., and K.J.C. conceived and designed the experiments. K.L.C., S.A.P., L.O., A.K., and D.H. performed the experiments and analyzed the data. K.L.C. and S.A.P. wrote the initial draft of the paper and outlined the figures. All authors reviewed and revised the manuscript.

